# Intravenous sulforhodamine B reduces alveolar surface tension, improves oxygenation and reduces ventilation injury in a respiratory distress model

**DOI:** 10.1101/2020.04.08.031435

**Authors:** You Wu, Tam L. Ngyuen, Carrie E. Perlman

**Author notes:** **Correspondence to:** Carrie E. Perlman, Ph.D., Stevens Institute of Technology, Department of Biomedical Engineering Castle Point on Hudson, Hoboken, NJ 07030.

## Abstract

In the neonatal (NRDS) and acute (ARDS) respiratory distress syndromes, mechanical ventilation supports gas exchange but can cause ventilation-induced lung injury (VILI) that contributes to high mortality. Further, surface tension, *T*, should be elevated and VILI is proportional to *T*. Surfactant therapy is effective in NRDS but not ARDS. Sulforhodamine B (SRB) is a potential alternative *T-*lowering therapeutic. In anesthetized male rats, we injure the lungs with 15 min of 42 ml/kg tidal volume, *V*_*T*_, and zero end-expiratory pressure ventilation. Then, over 4 hrs, we support the rats with protective ventilation – *V*_*T*_ of 6 ml/kg with positive end-expiratory pressure. At the start of the support period, we administer intravenous non-*T*-altering fluorescein (targeting 27 μM in plasma) without or with therapeutic SRB (10 nM). Throughout the support period, we increase inspired oxygen fraction, as necessary, to maintain >90% arterial oxygen saturation. At the end of the support period we sacrifice the rat; sample systemic venous blood for injury marker ELISAs; excise the lungs; combine confocal microscopy and servo-nulling pressure measurement to determine *T in situ* in the lungs; image fluorescein in alveolar liquid to assess local permeability; and determine lavage protein content and wet-to-dry ratio (W/D), both to assess global permeability. Lungs exhibit focal injury. Surface tension is elevated 72% throughout control lungs and in uninjured regions of SRB-treated lungs, but normal in injured regions of treated lungs. Sulforhodamine B administration improves oxygenation, reduces W/D and reduces plasma injury markers. Intravenous SRB holds promise as a therapy for respiratory distress.

**New and Noteworthy:** Sulforhodmaine B lowers *T*. Given the problematic intratracheal delivery of surfactant therapy for ARDS, intravenous SRB might constitute an alternative therapeutic. In a lung injury model, we find that intravenously administered SRB crosses the injured alveolar-capillary barrier, reduces *T* specifically in injured lung regions, improves oxygenation and reduces the degree of further lung injury. Intravenous SRB administration might help respiratory distress patients, including those with the novel coronavirus, avoid mechanical ventilation or, once ventilated, survive.

## Introduction

Neonatal (NRDS) and acute (ARDS) respiratory distress syndromes are high-incidence conditions (6, 8, 13, 42, 43, 57), and ARDS incidence has surely increased during the novel coronavirus pandemic. Mechanical ventilation supports gas exchange in NRDS and ARDS but also causes ventilation-induced lung injury (VILI), thus contributes to high rates of morbidity and mortality (4, 6–8, 10, 26, 39, 40, 43, 50, 51, 57, 61).

Ventilation injury has been attributed to a heterogeneous pattern in underlying lung injury, thus heterogeneous regional mechanics, in NRDS/ARDS and has been suggested to be proportional to alveolar interfacial surface tension, *T* (15, 36, 55, 58). The following *T-*dependent mechanical mechanisms have been proposed to cause VILI. First, as injured regions are diminished in size or collapsed, lung inflation may, due to interdependence, injuriously over-expand previously healthy adjacent tissue and thus cause injured regions to grow larger over time (5, 9, 15, 33, 36, 37). As flooded or collapsed regions typically open at a threshold opening pressure, *P*_*O*_, that is proportional to *T* (12, 16, 29), adjacent tissue may be most extended just before airway pressure reaches *P*_*O*_ and injury may be proportional to *T* (36). Second, ventilation may cause atelectrauma in injured regions (32, 44). And because re-opening stress should be proportional to *T*, injury may be proportional to *T* (2). Third, in local regions with heterogeneous alveolar flooding, we find that inflation injuriously exacerbates *T-* dependent stress concentrations in regional septa and that reducing *T* reduces inflation-induced over-extension injury (29, 36, 58). These mechanisms may co-exist.

Surface tension is elevated in NRDS, due to insufficient surfactant in the immature lungs, and believed to be elevated in ARDS, for various proposed reasons – plasma proteins, cholesterol or cell debris in edema liquid, upregulated phospholipase activity, oxidation of surfactant, aspirated acidic gastric contents and airway mucins accumulated by aspiration of gastric contents, with the greatest emphasis placed on plasma proteins (18–20, 22, 26, 31, 34, 35, 41, 46). In NRDS, exogenous surfactant therapy is highly effective (26). However mechanical ventilation can nonetheless injuriously increase alveolar-capillary barrier permeability, such that plasma proteins enter the airspace, and contribute to bronchopulmonary dysplasia (25, 26, 51). Further lowering *T* should further reduce VILI (58). In ARDS, unexpectedly, surfactant therapy has not reduced mortality (3). Surfactant dosage or delivery may be problematic (14, 27). An alternative means of lowering *T* in ARDS is required.

Previously, working in healthy isolated lungs, we found that alveolar administration of sulforhodamine B (SRB), a non-toxic dye approved as a food coloring in Japan (45, 56), lowers *T* up to 33% below normal (29). Sulforhodamine B is effective over a concentration range from 1 nM to 1 μM in the alveolar liquid, with greatest efficacy at 10 nM, and requires the presence of 3-12% albumin (29). The combination of SRB and albumin reduces *T* in the presence of the purported *T*-raising substances cell debris, hemoglobin and secretory phospholipase A_2_ and, as discussed previously, would likely be effective if *T* were elevated due to surfactant lipid oxidation (35). As SRB does not counter *T* elevation by acid or mucins, however, SRB would not likely be effective in the ∼12% of ARDS cases caused by gastric aspiration (13, 35). In heterogeneously flooded regions of isolated perfused lungs, alveolar SRB administration reduces both *T* and VILI (29).

In the present study, we test the ability of SRB, by lowering alveolar *T*, to reduce VILI in an *in vivo* lung injury model. Sulforhodamine B could potentially be administered by tracheal instillation, tracheal nebulization or intravenous injection. With airflow helping to drive distribution of solutions instilled or nebulized in the trachea (11, 14, 27, 52), delivery via the trachea to under-ventilated injured regions at appropriate concentration would be a significant challenge. Further, tracheal instillation of any liquid washes *T-* raising mucins to the alveoli, and SRB does not counter the effect of mucins (34, 35). Tracheal instillation is unlikely to be an effective route of SRB administration. Intravenous administration, however, should deliver SRB to the alveolar liquid in regions with elevated barrier permeability, at a concentration equal to or slightly less than that in the blood plasma. The elevated barrier permeability in respiratory distress also conveniently provides an appropriate albumin concentration in the alveolar liquid of injured regions. Thus, we administer SRB intravenously and target a plasma concentration of 10 nM, or 10x the minimum-effective alveolar concentration.

## Methods

We employ high tidal volume, *V*_*T*_, ventilation, an established means of generating a lung injury model (9, 47), in the first of with two distinct ventilation periods. We ventilate with high *V*_*T*_ for 15 min, to generate lung injury. Then we ventilate with a protective *V*_*T*_ for 4 hrs, to support the injured rat. We administer intravenous SRB after the injury period, at the start of the support period (Fig. 1), mimicking administration after the development of respiratory distress but prior to supportive treatment by mechanical ventilation. During the 4-hr support period, we track oxygenation. At the end of the support period, we sacrifice the rat, determine *T in situ* in the lungs and evaluate metrics of injury and inflammation.

**Figure 1:**
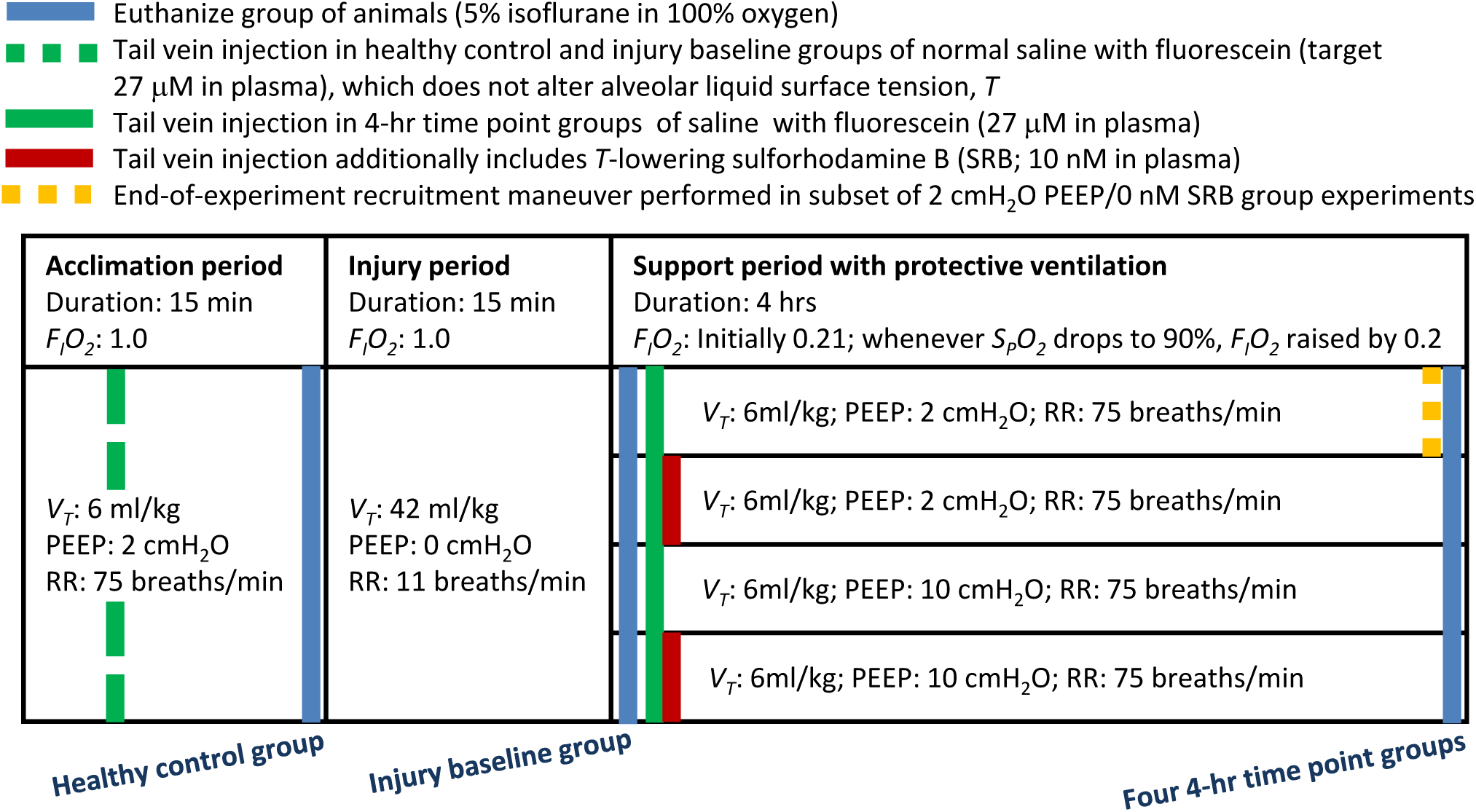
Protocol. Schematic of experimental protocol. PEEP: positive end-expiratory pressure; *F*_*I*_*O*_*2*_: fraction of inspired oxygen; *S*_*P*_*O*_*2*_: peripheral arterial oxygen saturation; *V*_*T*_: tidal volume; RR: respiratory rate. Euthanasia requires 20-40, 15-25 and 10-20 min in healthy-control, injury-baseline and 4-hr time-point groups, respectively.

### In vivo protocol

We handle all animals in accord with a protocol approved by the Stevens Institute of Technology Institutional Animal Care and Use Committee. We anesthetize male Wistar-Han rats (n=56, 270–375 g, Charles River, Wilmington, MA) with 3% isoflurane in 100% oxygen and perform a tracheotomy. We connect the tracheotomy tube to the circuit of a mechanical ventilator (model 683, 30 ml cylinder, Harvard Apparatus, Holliston, MA; compliance chamber between anesthesia vaporizer and ventilator inlet) with a stopcock that also connects a transducer for recording airway opening pressure, *P*_*AW*_.

#### Acclimation period

We ventilate the animal during a 15-min acclimation period with a fraction of inspired oxygen, *F*_*I*_*O*_*2*_, of 1.0, a nominal *V*_*T*_ of 6 ml/kg, a respiratory rate (RR) of 75 breaths/min (bpm) and a positive end-expiratory pressure (PEEP) of 2 cmH_2_O. We maintain body temperature at 37 °C and, every five minutes, record end-tidal CO_2_, *E*_*T*_*CO*_*2*_, with a capnograph and peripheral arterial oxygen saturation, *S*_*P*_*O*_*2*_, and heart rate (HR) with a pulse oximeter (PhysioSuite, Kent Scientific, Torrington, CT).

#### Injurious ventilation period

We injure the lungs by ventilating for 15 min with a *F*_*I*_*O*_*2*_ of 1.0, a *V*_*T*_ of 42 ml/kg, a RR of 11 bpm and zero end-expiratory pressure (ZEEP) (47). The ZEEP is achieved with a set PEEP of 2 cmH_2_O plus capnograph suction during the prolonged end-expiratory period.

#### Treatment and supportive ventilation period

We support the injured rat with protective ventilation: a *F*_*I*_*O*_*2*_ of 0.21, a *V*_*T*_ of 6 ml/kg, a RR of 75 bpm and a PEEP of 2 or 10 cmH_2_O. The start of support period is the injury-baseline time point. We administer a tail vein injection (normal saline base, total volume 100 μL) without (control) or with *T-*lowering SRB (target 10 nM in plasma; 341738, Sigma Aldrich, St. Louis, MO). In the injections for both control and experimental groups, we also include non-*T*-altering (28) fluorescein (target 27 μM in plasma; Cole Parmer, Vernon Hills, IL) for visualization and barrier permeability quantification under a fluorescent confocal microscope once the lungs are later excised. Calculating required dosages using a reported 45% hematocrit (Charles River, Wilmington, MA) and a rat-specific correlation between blood volume and body weight (30), we administer 299 μg/kg fluorescein without (control) or with 186 ng/kg SRB. We administer the injection 7.5 ± 3.5 min after the injury-baseline time point. From injury baseline, we ventilate for 4 hrs. Whenever *S*_*P*_*O*_*2*_ drops to 90%, we increase *F*_*I*_*O*_*2*_ by 0.20. If increasing *F*_*I*_*O*_*2*_ does not increase *S*_*P*_*O*_*2*_ above 90% within 5 min, we again increase *F*_*I*_*O*_*2*_ by 0.20. (Preliminary testing demonstrated that a *F*_*I*_*O*_*2*_ step size of 0.1 was often insufficient to increase *S*_*P*_*O*_*2*_ above 90% within 5 min in this model.) At the end of the 4-hr support period, we euthanize with 5% isoflurane in 100% oxygen.

#### Additional experimental groups

We asses two early-time-point control groups. In a healthy-control group, we anesthetize rats, begin acclimation period ventilation, administer a tail vein injection of fluorescein during the acclimation period and euthanize at the end of the acclimation period. In an injury-baseline group, we do the same but continue ventilating through the injury period before returning to protective ventilation with a PEEP of 2 cmH_2_O and euthanizing.

To investigate the cause of elevated *T* in our model, we also assess a recruitment-maneuver control group. We repeat the full protocol of the 2-cmH_2_O PEEP/0-nM SRB group and add, at the 4-hr time point, just prior to euthanasia, a single recruitment maneuver (RM) that raises peak inspiratory pressure (PIP), over 3-5 breaths, to 32-36 cmH_2_O.

### Post-mortem protocol

#### Thoracotomy and blood collection

After euthanasia, we close the tracheal stopcock with *P*_*AW*_ equal to PEEP and open the chest. We ligate the inferior vena cava and, proximal to the ligature, collect ∼4 ml blood, which we heparinize (20 U/ml; Penn Veterinary Supply Inc., Lancaster, PA) and centrifuge at 1000 x g and 4 °C for 10 min. We aliquot the supernatant and store the aliquots at -80 °C.

#### Lung isolation and imaging

We isolate the lungs and heart, position the lungs costal surface upward, photograph the lungs and place the lungs on the stage of an upright confocal microscope (SP5, Leica Microsystems, Buffalo Grove, IL). We image an injured surface region by bright-field and fluorescent confocal microscopy (x20, 0.5 N.A. water immersion objective; cover slip just contacting lung surface (60); for fluorescein fluorescence, 488 nm/≥493 nm excitation/emission). Then, we connect a house air source to the tracheal stopcock, open the stopcock to the house air and sequentially increase transpulmonary pressure, *P*_*L*_, to 30 cmH_2_O and decrease *P*_*L*_ to 15 cmH_2_O. In 4-hr time-point lungs, to quantify fluorescein intensity in alveolar flooding liquid, we obtain confocal images (x40, 0.8 N.A. water immersion objective; cover slip just contacting lung surface; 488 nm/≥493 nm excitation/emission; 798 gain) 20 μm below the pleural surface in three flooded regions, or fewer regions in the 40% of lungs in which we cannot locate three.

#### T determination

After identifying a surface alveolus in which to determine *T*, we subject the lungs to a regular volume history by cycling *P*_*L*_ twice between 5 and 15 cmH_2_O. We ensure that the size of the selected alveolus is responsive to pressure change, indicating that the airways leading to the alveolus are patent. We then hold *P*_*L*_ at 15 cmH_2_O and determine alveolar air pressure with the transducer at the trachea. We use servo-nulling pressure measurement (Vista Electronics, Ramona, CA) to determine liquid phase pressure in a surface alveolus. We use confocal microscopy to determine the three-dimensional radius of curvature of the air-liquid interface in the alveolus. And we calculate *T* from the Laplace relation (28, 34). In healthy-control and baseline-injury lungs, we determine *T* in the corner of an aerated alveolus. In 4-hr time-point lungs, we determine *T* in the corner of an aerated alveolus and at the meniscus (28) in a flooded alveolus.

#### Wet-to-dry ratio

We reduce *P*_*L*_ to 5 cmH_2_O, apply a vascular clamp (S&T B1-V, Fine Science Tools, Foster City, CA) between the right middle and caudal lobes and separate the caudal lobe distal to the clamp. We weigh the caudal lobe before and after drying at 76 °C for 24 hrs, and determine wet-to-dry ratio (W/D).

#### Lavage

We ligate the right middle lobe bronchus, cut the bronchus distal to the ligation, cannulate the bronchus and lavage the lobe with 1.54 mL/kg body weight room-temperature normal saline. We collect 127 ± 33 μL lavage liquid, which we centrifuge at 1000 x g and 4 °C for 10 min. We aliquot the supernatant and store the aliquots at -80 °C.

#### Lung fixation and histology

With *P*_*AW*_ still held at 5 cmH_2_O, we fix the remainder of the lungs to observe alveolar flooding pattern at ∼end-expiratory lung volume. We ligate the trachea, submerge the lungs in normal saline and microwave the lungs for 4 min to >60 °C (59). We transfer the lungs to 4% paraformaldehyde for 48 hrs and then to 70% ethanol. We obtain two hematoxylin- and eosin-stained, 3 μm-thick sagittal sections through the left lung, located at ∼1/3 and ∼2/3 the distance along the transverse axis (Histowiz, Brooklyn, NY).

#### Experimental series

We follow the same *in vivo* protocol in all experiments but vary the post-mortem protocol between series of experiments. We run separate experimental series for *T* determination in aerated alveoli and for microscopic imaging of injured lung regions at PEEP, and in those series perform select additional *post-mortem* procedures.

### Post-experimental analysis

#### Lung mechanics

We assess mechanics by analyzing the difference between PIP and PEEP which, for fixed PEEP and *V*_*T*_, and despite lack of plateau pressure, suggests respiratory system elastance. We determine the injury-baseline PIP-PEEP difference by averaging the value over 10 ventilation cycles starting 2 min after *V*_*T*_ reduction to 6 ml/kg and determine the 4-hr time-point PIP-PEEP difference by averaging the value over the last 10 cycles before commencing euthanasia. Starting at injury baseline, we fit an exponential function to the PIP-PEEP difference to determine the time constant of elastance increase.

#### Image analysis

In each fluorescent image of a flooded region at 15 cmH_2_O, we quantify fluorescein intensity in the liquid of three flooded alveoli. We average all intensity measurements from a given lung.

#### Histologic analysis

Histologic analysis of perivascular and alveolar edema is performed by Dr. Jerrold Ward (Global VetPathology, Montgomery Village, MD).

#### Protein extravasation and plasma biomarker quantification

We thaw lavage liquid or blood plasma. We determine protein concentration of lavage liquid by Bradford assay (Thermo Fisher Scientific, Waltham, MA). We perform enzyme-linked immunosorbent assays (ELISAs) of plasma for the soluble receptor for advanced glycation end-products (sRAGE; Mybiosource, San Diego, CA), surfactant protein D (SP-D; Lifespan Biosciences, Seattle, WA), tumor necrosis factor α (TNF-α; R&D Systems, Minneapolis, MN), interleukin 6 (IL-6; Invitrogen, Carlsbad, CA), plasminogen activator inhibitor 1 (PAI-1; Lifespan Biosciences) and von Willbrand factor (vWF; Lifespan Biosciences). For all assays, we follow manufacturer instructions.

#### Statistical analysis

We perform statistical analysis on all data as follows. We perform a Shapiro-Wilk test for normality and, if the data fail the normality test, log-transform the data. We test differences between experimental time points and the effects of independent factors by multiple linear regression. We report results as mean ± standard deviation and accept significance at p < 0.05.

### Results

One rat in the 10-cmH_2_O PEEP/0-nM SRB group, in which lung injury was greatest, died 3 hrs into the 4-hr support period. Data from this outlier are omitted from the results. All other rats survived to the planned euthanasia time points.

#### In vivo physiology

There are no significant differences over time or between groups in HR, *S*_*P*_*O*_*2*_ or body temperature, which average 368 ± 33 beats/min, 94.1 ± 1.2% and 37.0 ± 0.3 °C, respectively. End-tidal CO_2_ does not differ between groups but differs between the injury and support periods, in which it averages 21.5 ± 3.2 and 38.8 ± 5.3 mmHg (p < 0.001), respectively. Within each group, the difference in *E*_*T*_*CO*_*2*_ between periods is likewise significant at p < 0.001.

The *F*_*I*_*O*_*2*_ required to maintain *S*_*p*_*O*_*2*_ >90% does not vary with PEEP but is reduced by SRB. Without SRB, required *F*_*I*_*O*_*2*_ at 4 hrs is always between 0.6 and 1.0. With SRB, required *F*_*I*_*O*_*2*_ at 4 hrs is always 0.4. Thus, SRB improves oxygenation and reduces within-group variability in oxygenation. These results are presented as *S*_*p*_*O*_*2*_/*F*_*I*_*O*_*2*_ vs. time (Fig. 2).

**Figure 2:**
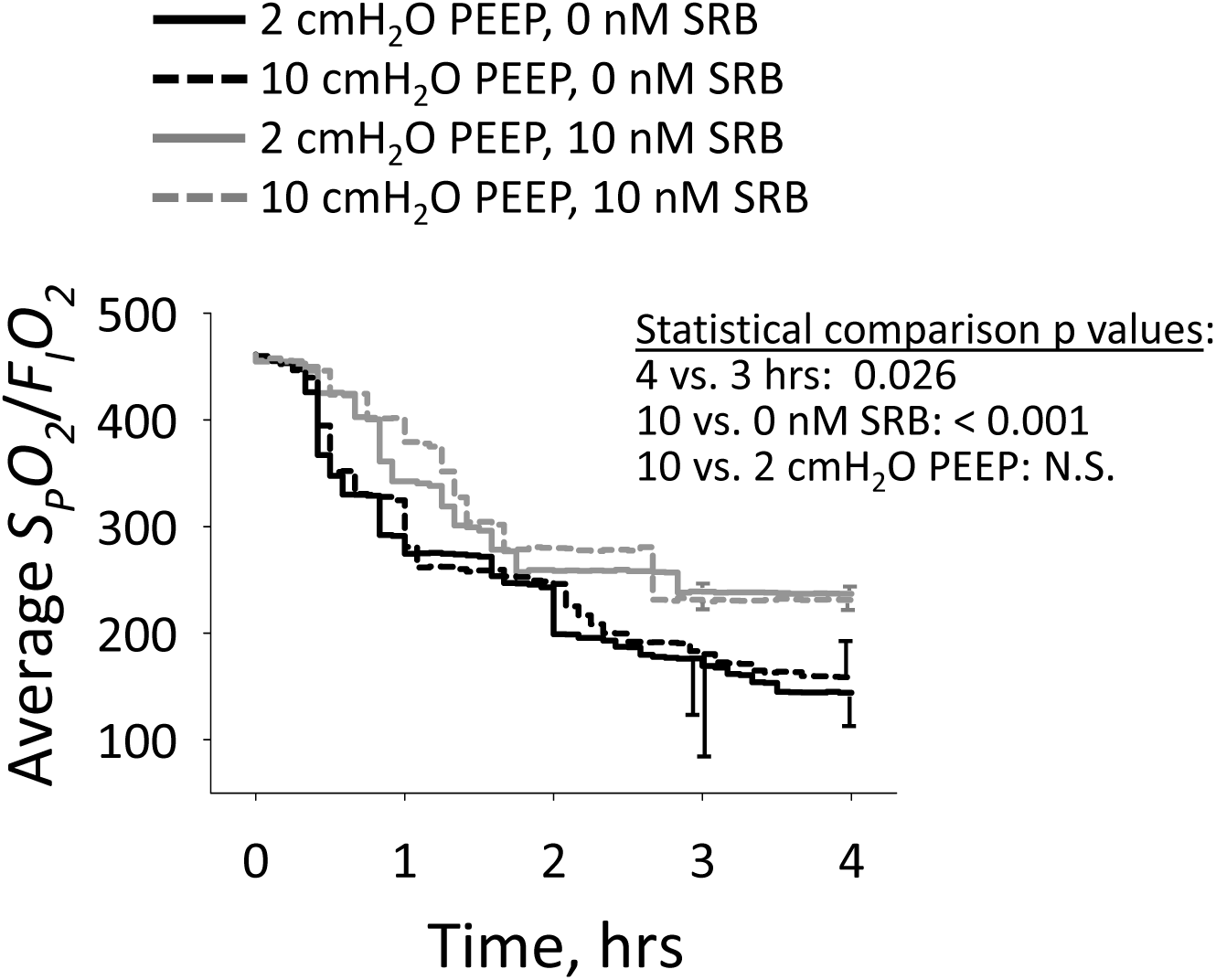
Oxygenation. Average *S*_*P*_*O*_*2*_/*F*_*I*_*O*_*2*_ vs. time, from injury baseline onward. n = 9-11/group. Statistical comparisons performed at 3- and 4-hr time points on log-transformed data.

#### In vivo lung mechanics

The anesthetic generally, but not always, prevents significant spontaneous breathing (>4 cmH_2_O decrease in minimum airway pressure below set PEEP). Significant spontaneous breathing during the acclimation period does not affect reported results. Significant spontaneous breathing later on affects one metric of barrier permeability, as detailed below, but not other metrics.

During the injury period, PIP averages 37 ± 2 cmH_2_O without a characteristic trend; PIP increases, decrease or remains constant in different animals. During the support period in the 2-cmH_2_O PEEP/0-nM SRB group, the PIP-PEEP difference increases thus elastance increases (Fig. 3). Administration of SRB increases the time constant of, thus delays, the increase in the PIP-PEEP difference. At injury baseline, the PIP-PEEP difference is greater with 10-than with 2-cmH_2_O PEEP. With 10-cmH_2_O PEEP, regardless of SRB administration, the PIP-PEEP difference does not change over the support period. At the 4-hr time point, the PIP-PEEP difference is the same regardless of PEEP or SRB administration.

**Figure 3:**
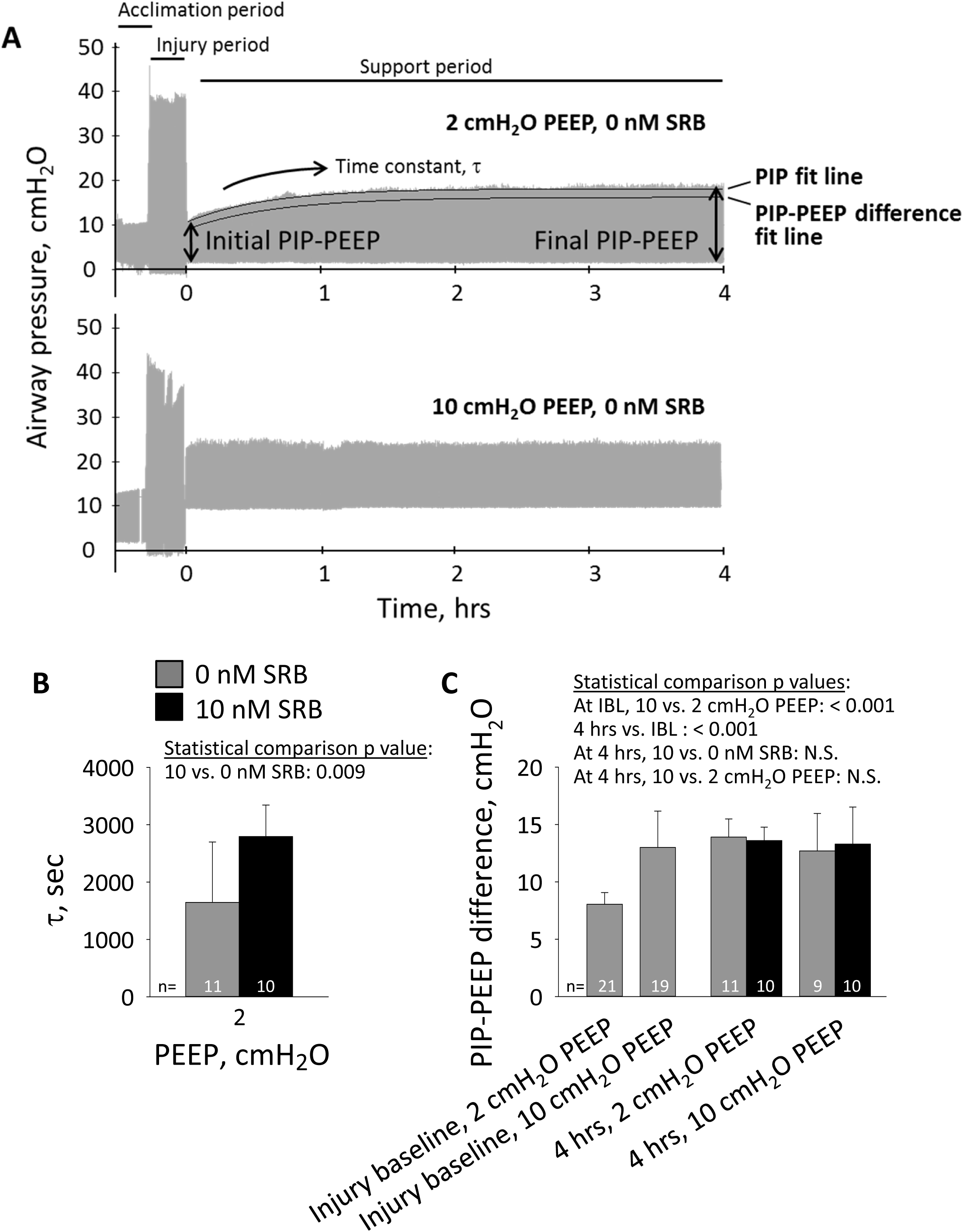
Lung elastance. **A**. Airway pressure traces, with indicated PEEP during support period, in absence of SRB administration. On 2-cmH_2_O PEEP trace, top black curve shows exponential fit to peak inspiratory pressure (PIP); bottom black curve shows exponential fit to difference between PIP and PEEP, which is shifted 2 cmH_2_O down from and has different time constant than top curve. With fixed PEEP and *V*_*T*_, PIP-PEEP difference indicates elastance. **B**. Time constant, τ, for increase in difference between PIP and PEEP, starting at injury baseline, in 2-cmH_2_O PEEP groups. C. Difference between PIP and PEEP at injury-baseline time point just prior to tail vein injection and at 4-hr time point at end of support period. For statistical comparisons, IBL: injury-baseline time point; 4 hrs: 4-hr time point; N.S.: not significant. At IBL, as SRB is not yet administered, 0- and 10-nM SRB groups are combined.

#### Lung appearance

All 4-hr time-point lungs exhibit focal hemorrhagic surface injury (Fig. 4), which has been observed previously in over-expansion induced lung injury (9). Neither of two blinded reviewers identified a difference in gross injury pattern due to PEEP or SRB administration. The same injury pattern is observed for 83% of injury-baseline and 50% of healthy-control lungs, perhaps due to euthanasia by isoflurane overdose in 100% oxygen (Fig. 1) (48, 49).

**Figure 4:**
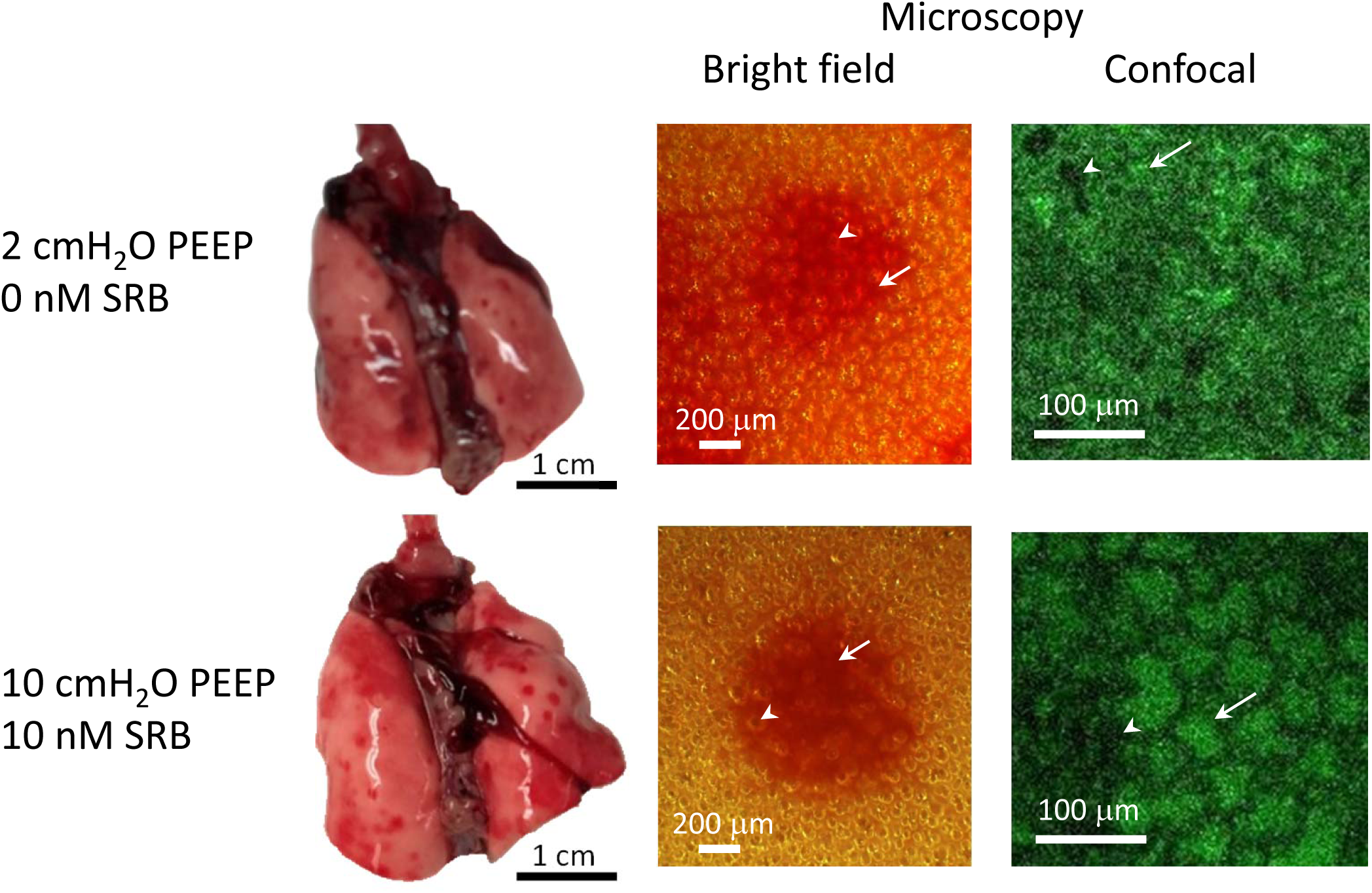
Lung appearance. Photographs show lungs isolated at 4-hr time point and at PEEP volumes, from indicated groups. Lungs exhibit focal hemorrhagic surface injury. Microscope images are of injured regions, with lungs still held at PEEP volume. Higher-magnification confocal images show fluorescein present in alveolar liquid and are located within central, injured regions of lower-magnification bright-field images. Arrows indicate shrunken, flooded alveoli (lack of air-liquid interface in bright-field image, green-labeled edema liquid in fluorescent image). Arrowheads indicate aerated alveoli. Color variation between bright-field images due to varying lamp illumination.

After lung isolation at PEEP and prior to lung inflation, bright-field microscopy of injured regions shows some alveoli to have disappeared (Fig. 4). As bright-field microscopy illuminates the alveolar air-liquid interface, the observed pattern indicates alveolar flooding or collapse. Fluorescent confocal microscopy of the same regions reveals that the alveoli that seem, by bright-field, to have disappeared are flooded and diminished. These findings are consistent across groups, from the 2-cmH_2_O PEEP/0-nM SRB group in which alveoli are least expanded to the 10-cmH_2_O PEEP/10-nM SRB group in which alveoli are most expanded. The presence of flooded/diminished alveoli on lung isolation at 10-cmH_2_O PEEP indicates that 10 cmH_2_O does not, in this model, maintain end-expiratory alveolar patency. Following increase of *P*_*L*_ to 30 cmH_2_O and decrease to 15 cmH_2_O, flooded alveoli remain small and aerated alveoli are over-expanded (Fig. 6C, below).

#### Surface tension

Surface tension at *P*_*L*_ of 15 cmH_2_O is normal (28) in healthy controls and not significantly elevated at injury baseline (Fig. 5). At 4 hrs, regardless of PEEP or SRB administration, *T* in aerated alveoli in uninjured regions is elevated. Also at 4 hrs, and regardless of PEEP, *T* in flooded alveoli in injured regions of control lungs is elevated to the same degree as *T* in aerated alveoli. Yet at 4 hrs, again regardless of PEEP, *T* in flooded alveoli of SRB-treated lungs is normal. Thus, intravenous SRB maintains low *T* specifically in injured regions. The maintenance of low *T* shows that both SRB and albumin, which must be present to facilitate SRB (29), are present in the liquid phase of injured regions. The region-specific reduction may be due, in part, to greater extravasation of 559 Da SRB (58), but is likely largely attributable to greater extravasation of 67 kDa albumin.

**Figure 5:**
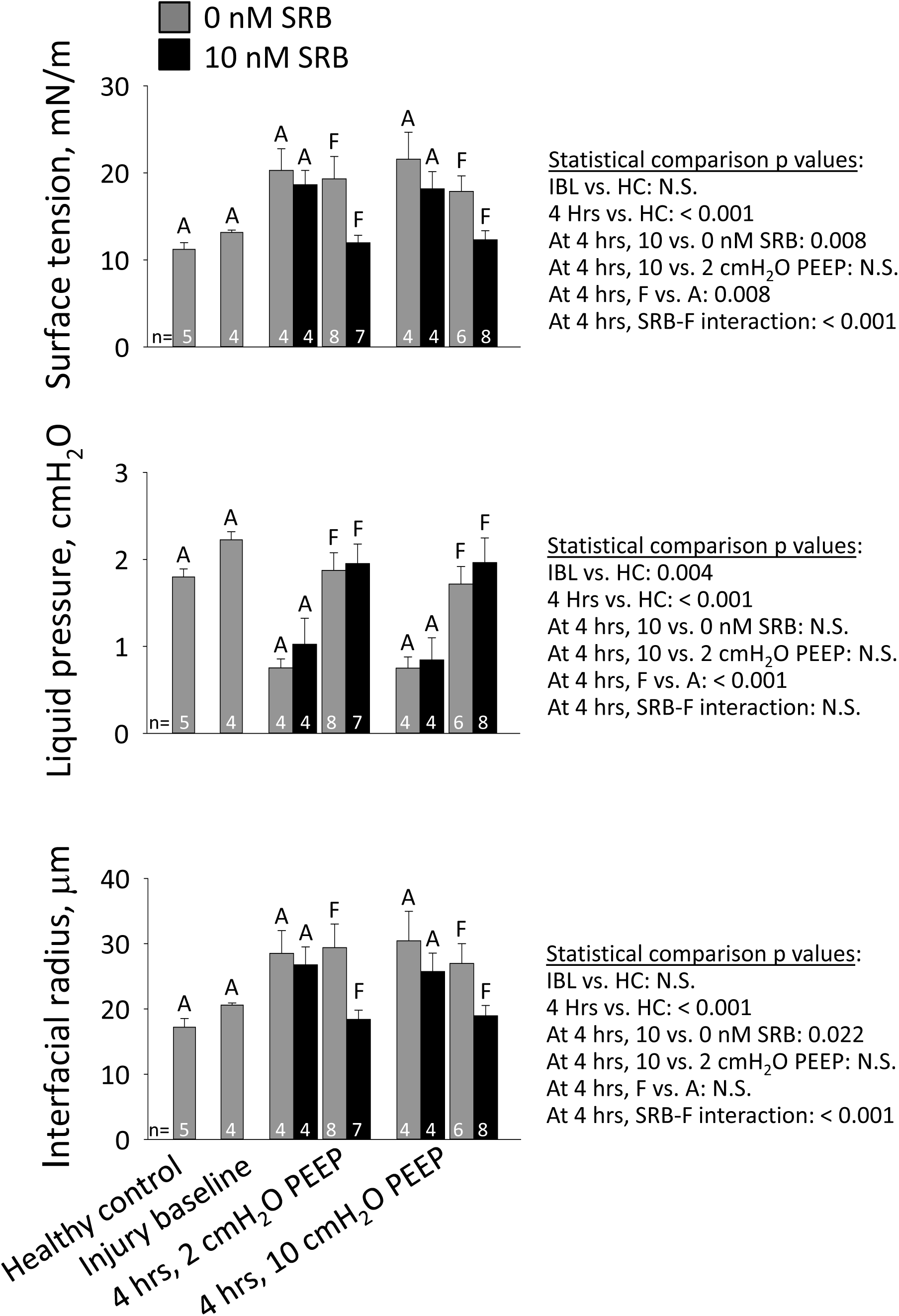
Surface tension and underlying measurements. Alveolar interfacial surface tension, T, is calculated from the Laplace relation after setting alveolar air pressure, *P*_*ALV*_, to 15 cmH_2_O; determining alveolar liquid phase pressure, *P*_*LIQ*_, in the liquid lining layer in the corner of an aerated (A) alveolus or below the meniscus of a flooded (F) alveolus by servo-nulling pressure measurement; and determining three-dimensional interfacial radius of curvature, *R*, from a *z*-stack of confocal microscopic images. For statistical comparisons, HC: healthy-control group.

In aerated alveoli, as expected from the Laplace relation, the trends in liquid pressure, *P*_*LIQ*_, and meniscus radius, *R*, are generally inversely and directly proportional, respectively, to that in *T* (Fig. 5). Nonetheless, the effect of *R* on *T* is greater than that of *P*_*LIQ*_. This difference is apparent from the results for the flooded alveoli. In untreated lungs, *T* is elevated in flooded alveoli principally due to an increase in *R* while *P*_*LIQ*_ is nearly normal. Similarly, SRB maintains normal, low *T* in flooded alveoli by maintaining a normal, small *R*. The small *R* with SRB treatment suggests the possibility of greater meniscus penetration into flooded alveoli, thus a thinner liquid layer.

With addition of a single recruitment maneuver at the 4-hr time point in 2-cmH_2_O PEEP/0-nM SRB experiments (Fig. 1, dashed orange line), the same focal injury pattern is observed as in other groups. The RM reduces *T* to the healthy-control level: 13.5 ± 0.5 mN/m (n=3) in aerated alveoli and 13.0 ± 1.4 mN/m (n=3) in flooded alveoli (for combined aerated and flooded alveoli, p < 0.001 vs. 2-cmH_2_O PEEP/0-nM SRB group without RM). These results suggest that *T* in our experiments without RMs is most likely elevated due to a low surfactant level. In uninjured aerated alveoli, that is almost certainly the case. In injured flooded alveoli, elevated *T* could alternatively be attributable to presence in the alveolar liquid of a *T*-raising substance that is masked by RM-induced secretion of additional surfactant. We cannot rule out the latter possibility.

#### Permeability and edema

We obtain global indications of permeability from W/D and lavage liquid protein content and a local indication of permeability from alveolar edema liquid fluorescein intensity. As the last metric is sensitive to spontaneous breathing, we exclude data for that metric from experiments with significant spontaneous breathing after the acclimation period.

Wet-to-dry ratio is elevated at 4 hrs (Fig. 6A). Administration of SRB reduces the increase in W/D over the support period. Lavage liquid protein concentration is likewise elevated at 4 hrs but, at that time point, does not differ between groups (Fig. 6B). Edema liquid fluorescein intensity data are only available at the 4-hr time point; no differences between groups are detected (Fig. 6C).

**Figure 6:**
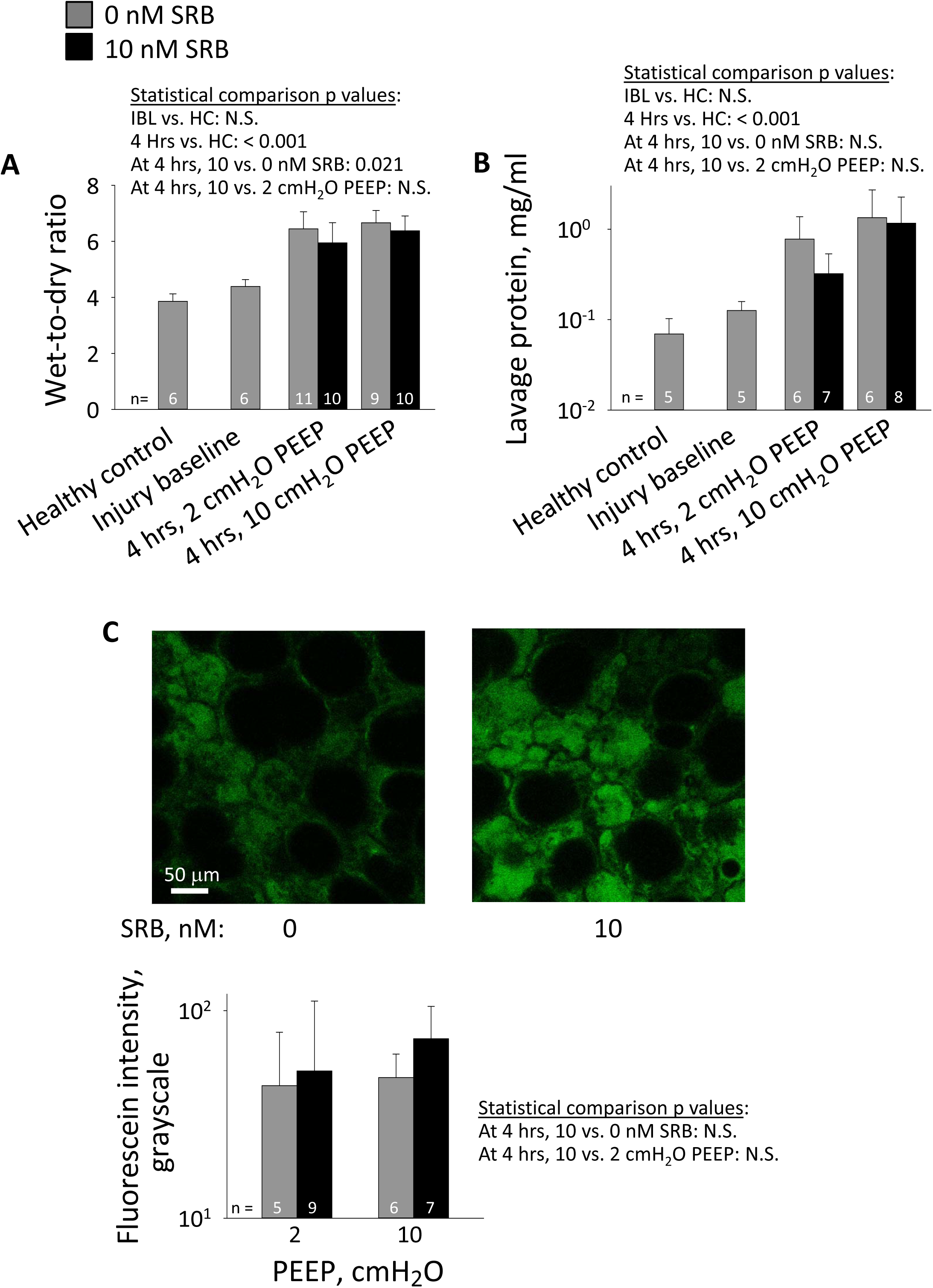
Permeability. **A**.Wet-to-dry ratio – a global permeability metric indicating combined injury severity and extent – of right caudal lobe. **B**. Total protein content of lavage liquid – an alternative global permeability metric – from middle right lobe. Statistical analysis for this metric performed on log-transformed data. **C**. Alveolar liquid fluorescein intensity – a local permeability metric specifically indicating injury severity. In 4-hr time-point lungs after inflation to 30 cmH_2_O and deflation to 15 cmH_2_O, confocal images obtained at 20-μm subpleural depth in flooded regions. Representative images from 10-cmH_2_O PEEP group exhibit average fluorescein intensities of 44 and 68 gray levels with 0- and 10-nM SRB, respectively. For display purposes, contrast of both images increased to same degree. Note typical pattern of diminished-sized flooded alveoli and over-expanded aerated alveoli. Graph presents group data. Statistical analysis for this metric performed on log-transformed data.

By histology (Fig. 7), high PEEP markedly worsens but SRB does not alter edema. Perivascular edema is more prevalent in the 4-hr time point/10-cmH_2_O PEEP groups than in all others. For alveolar edema, 1-3 foci are present in sections from injury baseline and 4-hr time point/2-cmH_2_O PEEP groups. One of the foci is always adjacent to the main airway branch point. More numerous foci are present in sections from 4-hr time point/10-cmH_2_O PEEP groups.

**Figure 7:**
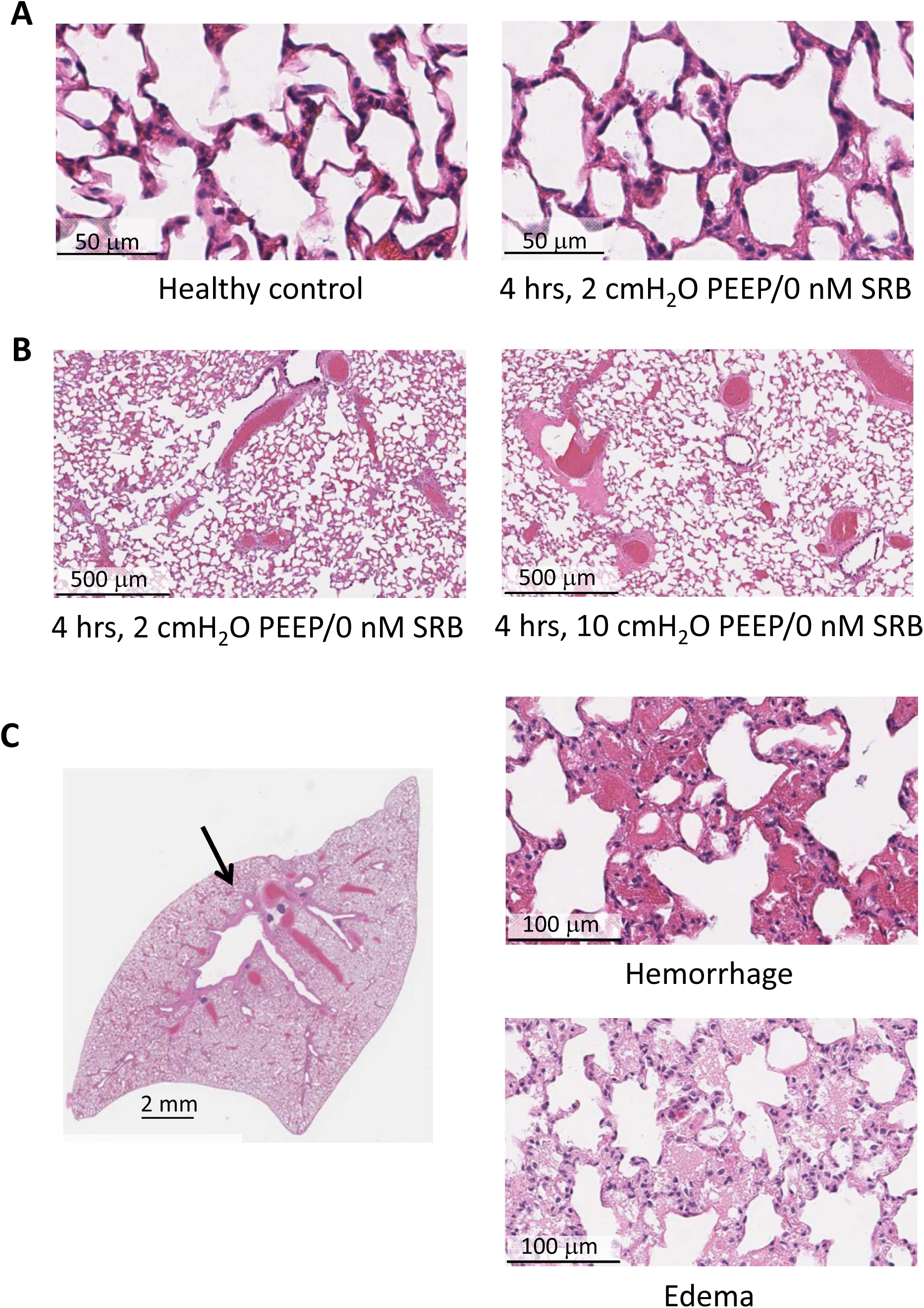
Histology. **A**.Representative images of uninjured regions in control and injured lungs. **B**. Perivascular edema is more frequent at high than low PEEP. **C**. Focal alveolar injury is always present adjacent to main airway branch point (arrow in left image of 2-cmH_2_O PEEP/0-nM SRB group lung). Injury is sometimes hemorrhagic (upper right image, from same lung as on left), sometimes edematous (lower right image, from 10-cmH_2_O PEEP/10-nM SRB group lung). Treatment with SRB does not alter histologic findings.

#### Injury markers

The high *V*_*T*_ period initiates injury and injury progresses during the 4-hr support period (Fig. 8). Treatment with SRB reduces levels of sRAGE, an indicator of alveolar epithelial type I cell injury; TNF-α, a cytokine; and PAI-1, an indicator of systemic inflammation. High PEEP ventilation increases levels of SP-D, an indicator of alveolar epithelial type II cell injury; IL-6, a cytokine; and PAI-1. Neither SRB nor PEEP affects levels of vWF, another indicator of systemic inflammation. Detection of lung injury markers in the plasma suggests a pronounced degree of lung injury.

**Figure 8:**
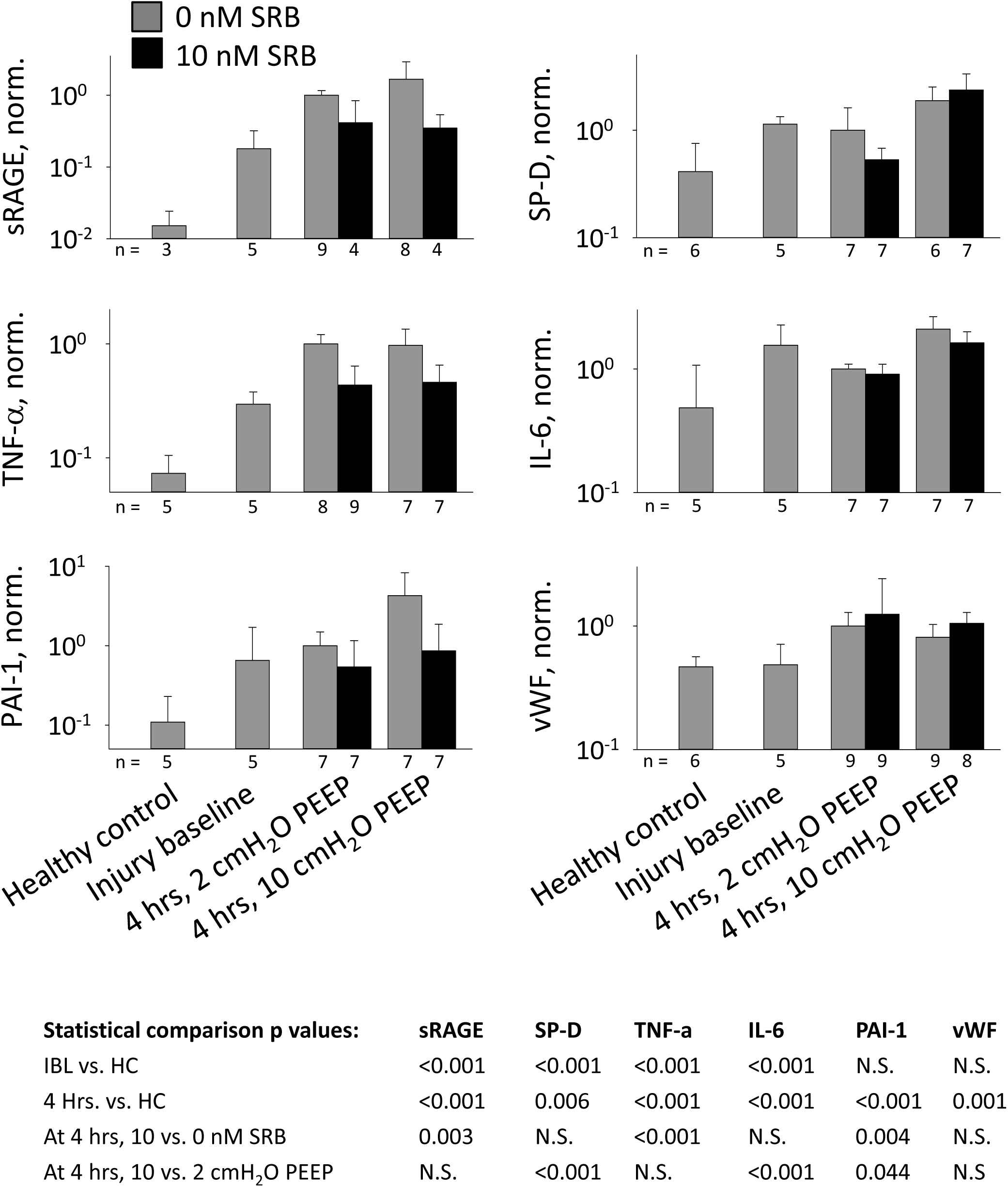
Injury markers. Injury markers quantified by enzyme-linked immunosorbent assays of blood plasma: soluble receptor for advanced glycation end products (sRAGE), surfactant protein D (SP-D), tumor necrosis factor α (TNF-α), interleukin 6 (IL-6), plasminogen activator inhibitor 1 (PAI-1) and von Willbrand factor (vWF). Data normalized by average value of 2-cmH_2_O PEEP/0-nM SRB group. Statistical analysis for all injury markers performed on log-transformed data.

## Discussion

We injure the lungs with high *V*_*T*_ ventilation. The protocol causes focal hemorrhagic lung-surface injury. In injured regions of untreated controls, surface tension is elevated only in the absence of a recruitment maneuver. Thus, *T* is likely elevated due to a low surfactant level. In injured regions of SRB-treated rats, sufficient SRB and albumin diffuse from blood to alveolar liquid to maintain normal *T* in the absence of a RM. The SRB-plus-albumin-induced maintenance of normal *T* in the presence of a likely-reduced surfactant level, like SRB-plus-albumin-induced reduction of *T* below normal in healthy lungs (29), supports the concept that SRB-plus-albumin improves the efficacy of native lung surfactant.

Elevation of *T* in ARDS has been attributed to various causes, with the greatest emphasis placed on plasma proteins in edema liquid (18–20, 22, 31, 34, 35, 41, 46). We and others have shown (21, 23, 28, 35, 53) that blood plasma does not raise *T* in the presence of an intact surfactant layer. However, it has not been known whether alveolar diminishment or collapse in an injured lung would rupture the surfactant layer and thus permit plasma contents to adsorb and raise *T*. Our 2-cmH_2_O PEEP, 0-nM SRB recruitment-maneuver experiments in the present study address that question. That a RM normalizes *T* in non-SRB-treated injured alveoli flooded with proteinaceous edema liquid indicates that the surfactant layer remains intact even when alveolar size is markedly diminished at the end of expiration (Fig. 4, top row). Further, the efficacy of SRB in flooded, hemorrhagic regions (Fig. 5) shows that SRB is effective in the presence of whole-blood components.

Given that an intact surfactant layer prevents many impurities from raising *T* (21, 23, 28, 35, 53), co-adsorption of surfactant and impurities in *in vitro* surfactometers introduces an artifact into *in vitro T* determinations. It is thus notable that the present study is the first to test a *T*-lowering therapy in an *in vivo* lung injury model and then to determine *T* directly in the lungs without disrupting the *in situ* surfactant layer.

The present study is also the first to test intravenous administration of a *T-*lowering therapy. While lung-specific administration of a pulmonary therapeutic holds appeal, effective tracheal administration of SRB would be challenging for reasons discussed above. We show here that SRB administration at appropriate concentration to injured lung regions is easily accomplished via the vasculature (Fig. 5). The ease of intravenous access in a hospital setting adds to the appeal of this route of administration.

It has been observed that injured regions grow over time (5, 9), indicating that ventilation stresses initially-healthy tissue adjacent to injured regions and eventually injures that tissue. This observation corresponds to the first of three possible mechanical mechanisms of VILI outlined in the Introduction. Our results suggest that SRB may interrupt that VILI mechanism – SRB may, at the expense of exacerbating existing injury, retard the growth of injured regions. We find that SRB reduces W/D and tends to reduce lavage protein concentration, both global permeability metrics indicating combined severity and spatial extent of injury, but tends to increase edema liquid fluorescein intensity, a local permeability metric specifically indicating injury severity (Fig. 6). Sulforhodamine B creates a spatial *T* differential – lower *T* in injured than adjacent healthy regions (Fig. 5) – thus should increase the maximal stress applied to injured tissue and reduce the maximal stress applied to adjacent healthy tissue. Maximal stress may be applied at peak inflation or, if injured regions cyclically collapse and re-open, just before *P*_*AW*_ reaches *P*_*O*_. In the latter case SRB, by lowering *P*_*O*_, should further reduce peak stress applied to adjacent healthy tissue from that just below the original, higher *P*_*O*_ to that below the new, lower *P*_*O*_. As the process of injured region growth is one of positive feedback, the small SRB-induced reduction in W/D observed over 4 hrs in this study might become more pronounced over a longer time. The SRB-induced reduction in markers of injury and inflammation (Fig. 8) supports the concept that SRB reduces overall VILI in the lungs. Although it is not known whether SRB has direct anti-inflammatory effects, SRB most likely acts by reducing *T* and, in turn, reducing ventilation-induced mechanical injury that is upstream of ventilatory exacerbation of inflammation. Given that the two clinically effective interventions for VILI are mechanical (4, 17), SRB action via an upstream mechanical mechanism suggests potential for clinical translation.

Sulforhodamine B administration improves oxygenation (Fig. 2), perhaps via multiple mechanisms. The SRB-induced reduction in W/D should contribute to the improved oxygenation, whether the reduction in W/D is due to reduced spatial extent of injury or reduced flooding in injured regions. The possibility that SRB may reduce flooding severity in injured regions is supported by the SRB-induced decrease in interfacial radius in flooded alveoli (Fig. 5), which suggests a thinner liquid layer. Sulforhodamine B-induced low *T* may additionally improve oxygenation by reducing *P*_*O*_ of injured alveoli, thus recruiting gas exchange area at lower volume; if the liquid layer is thinned in injured regions once opened, by allowing capillaries to expand (24, 54) and thus, again, increasing gas exchange area; or by increasing compliance of injured regions, thus reducing inter-regional differences in time constants and reducing ventilation-perfusion mismatch.

Comparing the effects of *T* and PEEP in control (0-nM SRB) lungs elucidates relative contributions of *T* and tissue elastance to respiratory system elastance. At injury baseline, the PIP-PEEP difference is greater with 10- than 2-cmH_2_O PEEP (Fig. 3C) due to increased *T* and tissue elastance at increased lung volume. Over the next 4 hrs, *T* increases throughout control lungs in both PEEP groups (Fig. 5). (With *T* always determined at *P*_*L*_ of 15 cmH_2_O, Fig. 5 shows that *T* increases over time in both the 2- and 10-cmH_2_O PEEP groups but does not show actual *T* in either group during support period ventilation. During that period, because PEEP increases lung volume and thus *T, T* would be higher in the 10- than 2-cmH_2_O PEEP group.) In contrast to the time-dependent increase in *T*, which is consistent across PEEP groups, there is only a time-dependent increase in PIP-PEEP difference in the 2-, not 10-, cmH_2_O PEEP group (Fig. 3). Consistent with a prior finding of ours (38), these results suggest that *T* may only affect lung mechanics at low lung volumes.

Higher PEEP is insufficient to maintain end-expiratory alveolar patency in this model (Fig. 4, bottom row), thus has a generally detrimental effect. Higher PEEP fails to improve oxygenation (Fig. 2) and increases multiple plasma injury markers (Fig. 8).

The SRB-induced reduction in *T* affects select metrics. Sulforhodamine B reduces permeability as indicated by W/D, improves oxygenation, delays increase of elastance and reduces several plasma injury markers, but does not alter other metrics of permeability, final lung elastance or histologic lung appearance. The discrepancies between these results may be attributable to the varying sensitivities of the different measurements. Most notably, due to the low solubility of oxygen in water, oxygen diffusion through edema liquid is highly sensitive to liquid thickness. It is not surprising that a small reduction in W/D would improve oxygenation before causing apparent changes in lung compliance or histologic appearance.

The animal model of this study has relevance to both ARDS and NRDS. We performed the study in adult rats with mature lungs such as those in which ARDS develops. The spatially heterogeneous ventilation injury and proteinaceous edema liquid are common to ARDS and NRDS. The cause of elevated *T* is likely a low surfactant level, which is that of NRDS. Nonetheless, SRB holds potential for lowering *T* in both ARDS and NRDS. We showed in an isolated rat lung model that SRB is likely to reduce *T* under ARDS conditions (35). And as SRB improves the efficacy of surfactant, SRB might be considered as a complement or alternative to surfactant therapy in NRDS.

Our study, however, has limitations. With a compliance chamber between anesthesia vaporizer and ventilator, pressure at the ventilator inlet may have been positive such that actual *V*_*T*_ may have exceeded nominal *V*_*T*_. Nonetheless, consistent injury-period PIP levels indicate consistent injury across animals. Without use of a paralytic, spontaneous breathing affects alveolar liquid fluorescein intensity results. As no means exists for determining *T in vivo*, we determine *T* in lungs isolated following animal sacrifice. We do not know if the focal injury pattern present on the lung surface is also present in the lung interior. With just two histologic sections from each lung, we may miss interior injury.

Alternatively, by increasing *P*_*L*_ to 30 cmH_2_O before fixation at 5 cmH_2_O, we may alter the alveolar collapse or flooding pattern from that at PEEP *in vivo*. Results for two of our permeability metrics do not differ significantly between 4-hr time-point groups, so our discussion of implications is speculative. Last but not least, we use only male rats. However, neither surface tension nor response to SRB appears to vary with gender (35).

In summary, we demonstrate that intravenous SRB administration lowers alveolar liquid *T* in regions with barrier injury. In this model of heterogeneous lung injury, intravenous SRB administration constitutes a novel approach to simultaneously improving oxygenation and reducing ventilation injury. As ventilation injury can occur with spontaneous breathing in injured lungs (1), intravenous SRB has the potential to help not-yet ventilated respiratory distress patients avoid mechanical ventilation as well as to help ventilated respiratory distress patients survive.

## Acknowledgements

We are grateful to Dr. Michelle Gong of Montefiore Medical Center for her input on experimental design and to Dr. Joseph Lucas of Vital Statistics, Inc. for his guidance on statistical analysis.

